# mRNA interactions promote cotranslational association of heteromeric membrane proteins

**DOI:** 10.1101/2025.07.01.662506

**Authors:** Lisandra Flores-Aldama, Annabelle S. Hoth, Lucas de Carvalho, Ian Seim, Fang Liu, Amy S. Gladfelter, Gail A. Robertson

## Abstract

Heteromeric membrane proteins play crucial physiological roles, yet how they are formed remains poorly understood. Heteromeric hERG1a/1b ion channels, essential for maintaining normal cardiac rhythm, assemble via cotranslational association of their encoding mRNAs. We hypothesized that direct *hERG1a* and *1b* mRNA interactions facilitate this process. Using fluorescence colocalization and free energy of binding predictions, we found that *hERG1a* and *1b* mRNAs form specific heterotypic condensates *in vitro*, suggesting direct interactions. When *hERG1a* mRNA was altered by synonymous mutations predicted to reduce its structural diversity and ability to interact with other mRNAs, overlap with *hERG1b* was dramatically diminished both *in vitro* and in cells, indicating weakened interactions. Reducing *hERG1a* structural diversity also influenced its translational complexes, defined by overlapping between fluorescently labeled mRNA and encoded protein (centroids within 400 nm). Whereas most of the wild-type *hERG1a* mRNA translates within heterotypic complexes, likely reflecting the biogenesis of hERG1a/1b heteromeric assemblies, reducing *hERG1a* structural diversity yielded more homotypic translational condensates and fewer hERG1a/1b heterotypic ones. This result suggests that the strength of mRNA interactions impact ion channel biogenesis. Further analysis of the heterotypic translational complexes revealed two distinct classes: a) simultaneous translation of both subunits and b) sequential association of fully translated hERG1b with translating hERG1a. Notably, reducing *hERG1a* structural diversity and interactions with *1b* shifted translation toward the sequential mode. These findings identify a new role of mRNA sequence, structure, and interactions in orchestrating the cotranslational association of important heteromeric membrane proteins.

**SIGNIFICANCE:** The physiological diversity of membrane protein complexes arises in part from the mixing of related yet functionally distinct subunits such as those forming “heteromeric” ion channels. How this mixing is achieved is not well understood. Here, we show that subunits composing heteromeric hERG1a/1b ion channels, crucial for normal cardiac function, predominantly associate in complexes where both subunits are simultaneously translated. This cotranslation is promoted by the association of the encoding mRNAs. Silent mutations disrupting *hERG1a* and *1b* mRNA association, without altering the encoded protein, diminish simultaneous translation in favor of hERG1a translational complexes alone or in association with fully synthesized hERG1b. This study highlights the significant impact of RNA sequence, structure, and association on the formation of heteromeric membrane proteins.

## INTRODUCTION

Numerous cellular processes are carried out by heteromeric protein complexes. Transmembrane receptors, transporters, and ion channels mediating chemical and electrical signaling are frequently heteromers of subunits of the same or distinct gene families (1–8). Because the structural, biophysical, and pharmacological properties of protein complexes depend on their subunit composition, heteromerization is proposed to enhance protein functional heterogeneity and specify their physiological roles (1–5, 9–13). Despite many studies describing the relation between heteromeric ion channel composition and functional properties, the underlying assembly mechanism is not well understood.

Heteromeric ion channels conduct multiple ionic currents that are essential for normal cardiac function (14–17). For example, the I_Kr_ current, which is critical for repolarizing the ventricular action potential, is conducted by heteromeric *ether-à-go-go* related gene (ERG) ion channels (4, 14, 18, 19). In humans and rodents, two highly conserved subunits, ERG1a and 1b, are encoded by alternate transcripts of the *KCNH2* gene (4, 20). *ERG1a* mRNA comprises 15 exons, the first five of which are substituted by a single alternative exon in *ERG1b* (4). ERG1a and 1b subunits differ only in their amino (N-) termini, with ERG1a containing a *Per-Arnt-Sim* (PAS) domain and ERG1b a short and unique N-terminal region (4). In humans, mutations in either the hERG1a or hERG1b subunit are linked to type 2 long QT syndrome (LQT2), and their coordinated expression and heteromeric assembly are essential for maintaining normal cardiac excitability (10, 14, 21–25). Thus, the association between hERG1a and 1b subunits is fundamental to normal cardiac function.

Previous studies showed that the heteromerization of hERG1a/1b channels occurs during early biogenesis and is accompanied by the association of their encoding mRNAs during translation (26, 27). This association was observed in cardiomyocytes derived from human induced pluripotent stem cells (hiPSC-CMs) and reconstituted in HEK-293 cells co-expressing hERG1a and 1b subunits, suggesting the elements required for cotranslational assembly are conserved in multiple cell types (27). Nevertheless, the mechanism underlying the association of *hERG1a* and *1b* mRNAs remains unknown.

Here, we tested the hypothesis that direct interactions between *hERG1a* and *1b* mRNA promote the cotranslational assembly of their encoded subunits. Indeed, we found that fluorescently labeled *hERG1a* and *1b* mRNAs directly associated *in vitro* to form condensates. Their interaction was influenced by *normalized ensemble diversity* (NED), a thermodynamic parameter that quantifies the number and diversity of structural conformations an RNA molecule adopts at equilibrium (28, 29). Specifically, we found that reducing *hERG1a* NED, or structural diversity, weakened its association with *hERG1b*, both *in vitro* and in cells. Using colocalization between hERG1a and 1b mRNA and encoded proteins, we showed that biogenesis mainly occurs in heterotypic complexes where both subunits are simultaneously translated. In live-cell experiments, reduced *hERG1a* structural diversity led to more homotypic translational complexes. Notably, this change was accompanied by a shift in the heterotypic translational profile from simultaneous to sequential association of nascent hERG1a with fully translated hERG1b. These findings suggest that translational association of hERG1a and 1b subunits is governed, at least in part, by direct mRNA interactions shaped by the sequence-encoded structural diversity of the transcripts.

## RESULTS

### *In vitro* reconstitution shows direct and specific *hERG1a* and *1b* mRNA interaction

To investigate whether *hERG1a* and *1b* mRNAs directly interact to form heterotypic complexes, we studied their behavior in an *in vitro* system devoid of intracellular macromolecules that might bridge their association. Independently transcribed *in vitro*, *hERG1a* and *1b* mRNAs were fluorescently labeled with Cy3 and Cy5 fluorescent dyes, respectively, and mixed in a 1:1 ratio within a minimum system containing polyamines and cations known to stabilize RNA structures under physiological conditions (30–35) (Fig. 1A). To mimic their naturally low abundance in cells, we used nanomolar *hERG1a* and *1b* mRNA concentrations (36–38). Under these conditions, mRNAs spontaneously formed condensates whose size and number were qualitatively affected by changes in the ionic strength and mRNA concentration (*SI Appendix*, Fig. S1). This behavior is consistent with that previously reported for nucleic acid condensates reconstituted *in vitro* (39, 40).

**Figure 1.**
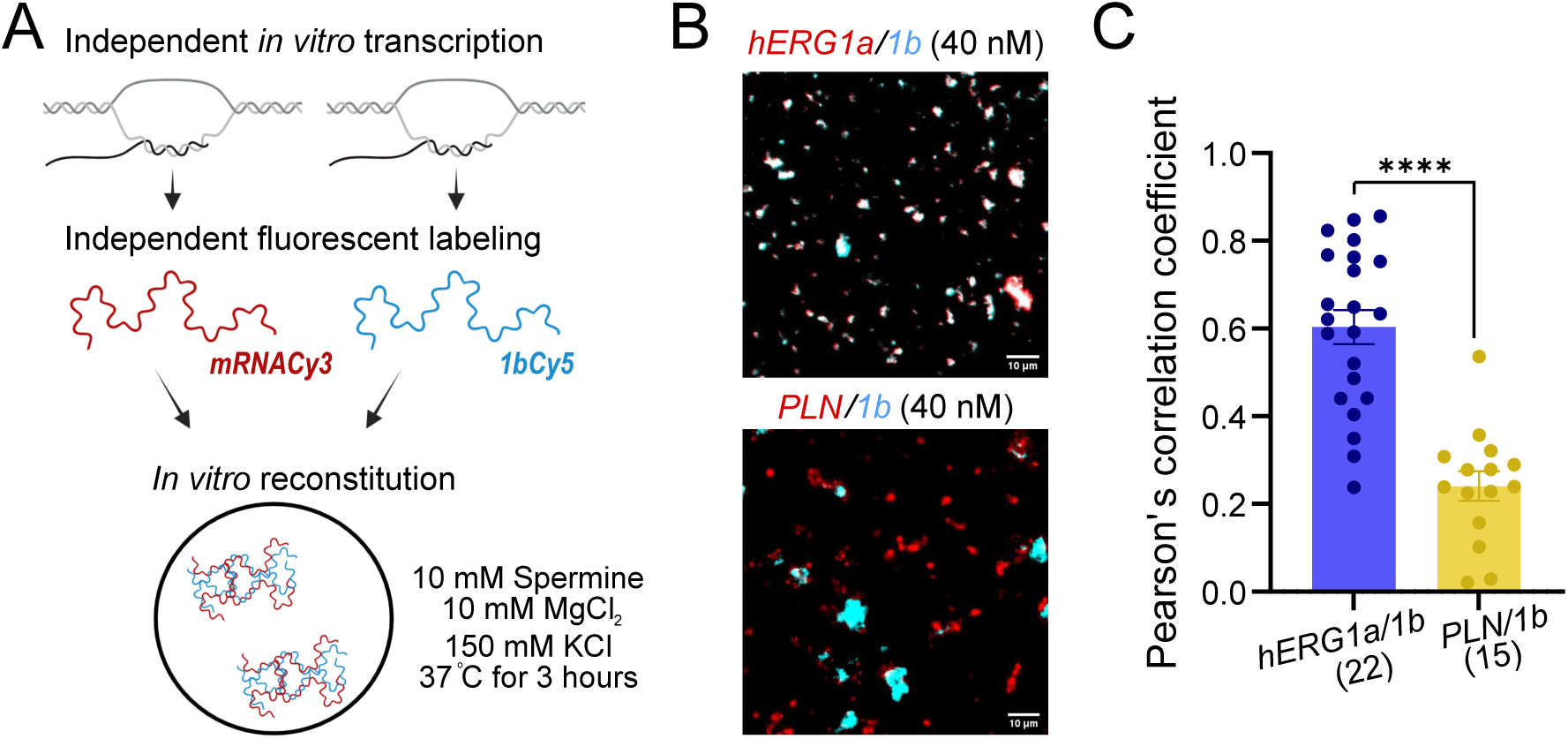
*In vitro* reconstitution shows direct and specific *hERG1a* and *1b* mRNA interaction. **A.** Schematics of the experimental design used for the *in vitro* reconstitution of heterotypic RNA mixes (*for more detail see Methods*). **B.** Representative images of *in vitro* reconstituted *hERG1a/1b* and *PLN/1b* mRNA pairs in a 1:1 ratio to 40 nM total RNA concentration in physiological K^+^ levels. The white signal indicates the overlap of spectrally separable fluorescent dyes. Scale bars represent 10 μm. **C.** Comparison of the Pearson’s correlation coefficient of *hERG1b* mRNA fluorescent signal overlap with *hERG1a* or *PLN* mRNAs. Bars represent mean + SEM. ****: p<0.0001. Single points indicate independent experiments and numbers in parenthesis represent replicates.

The colocalization of *hERG1a* and *1b* mRNAs was apparent from the overlap of their respective fluorescent signals, appearing as white and quantified by Pearson’s correlation coefficient (Fig. 1B, C; *see Methods*). The Pearson’s coefficient was 2.5 times greater than when *hERG1a* was replaced with a control mRNA encoding phospholamban (*PLN*), a functionally unrelated membrane protein also expressed in ventricular cardiomyocytes. Despite its similar length and NED to *hERG1a* and *1b* mRNAs (see below), the *PLN* transcript has a less favorable predicted interaction with *1b* (Table 1, *SI Appendix*, Fig. S2), making it a suitable control for nonspecific mRNA association. These observations indicate that *hERG1b* directly and specifically interacts with *hERG1a in vitro*.

**Table 1.** *Normalized Ensemble Diversity* (NED) for mRNAs used in the study. NED was calculated by dividing the ensemble diversity, predicted using RNAfold from the ViennaRNA package, by the length (in nucleotides) in each mRNA.

| mRNA | NED |
| --- | --- |
| *hERG1aWT* | 0.25 |
| *hERG1b* | 0.24 |
| *PLN* | 0.25 |
| *hERG1aLN* | 0.05 |
| *hERG1aSN* | 0.22 |
| *hERG1aHN* | 0.38 |
| *hERG1aWT-BFP* | 0.24 |
| *hERG1aLN-BFP* | 0.07 |

### *hERG1a* structural diversity influences its association with *1b in vitro*

Theoretically, *hERG1a* and *1b* mRNA structure and association depend on a dynamic equilibrium of non-covalent, intra- and intermolecular interactions (41–43). These interactions include complementary and non-canonical base pairing, base stacking, hydrogen bonds, London-Van der Waals, and electrostatics, all influenced by the surrounding microenvironment conditions and RNA physicochemical properties such as GC content and nucleotide sequence and length (28, 30–35, 41, 42, 44). Accordingly, each mRNA molecule can adopt an ensemble of structures based on these dynamic interactions (29). This property is embodied quantitatively by the *normalized ensemble diversity* (NED), defined as the average number of contiguous base pairs that must be rearranged for a given RNA molecule to transition from one thermodynamically favorable conformation to another, normalized by its length (28, 29, 45). RNAs with low NED are characterized by stable local structural motifs with high intramolecular base pairing probability (28, 29, 42). RNAs with higher NED present more opportunities for conformational changes and potential interactions with other RNA molecules.

To determine whether mRNA conformational heterogeneity can influence the interaction between *hERG1a* and *1b* mRNAs, we manipulated *hERG1a* NED using a novel genetic algorithm developed for this purpose (*see Methods*; (46)). We generated synonymously mutated *hERG1a* mRNAs with varying intramolecular base pairing probability, predicted difference of free energy between unfolded and folded states, and structural diversity, resulting in lower (*1aLN*), similar (*1aSN*), or higher (*1aHN*) NED than *hERG1aWT* (Fig. 2A, Table 1). Mutations were introduced only in the coding sequence without affecting the UTRs (*SI Appendix*, Supplementary File 1). Wild-type and mutant *hERG1a* mRNAs encoded appropriately sized hERG1a proteins that are comparably expressed, as shown by western blot analyses with antibodies targeting hERG1a Nterminal or pan-hERG1 regions (Fig. 2B).

**Figure 2.**
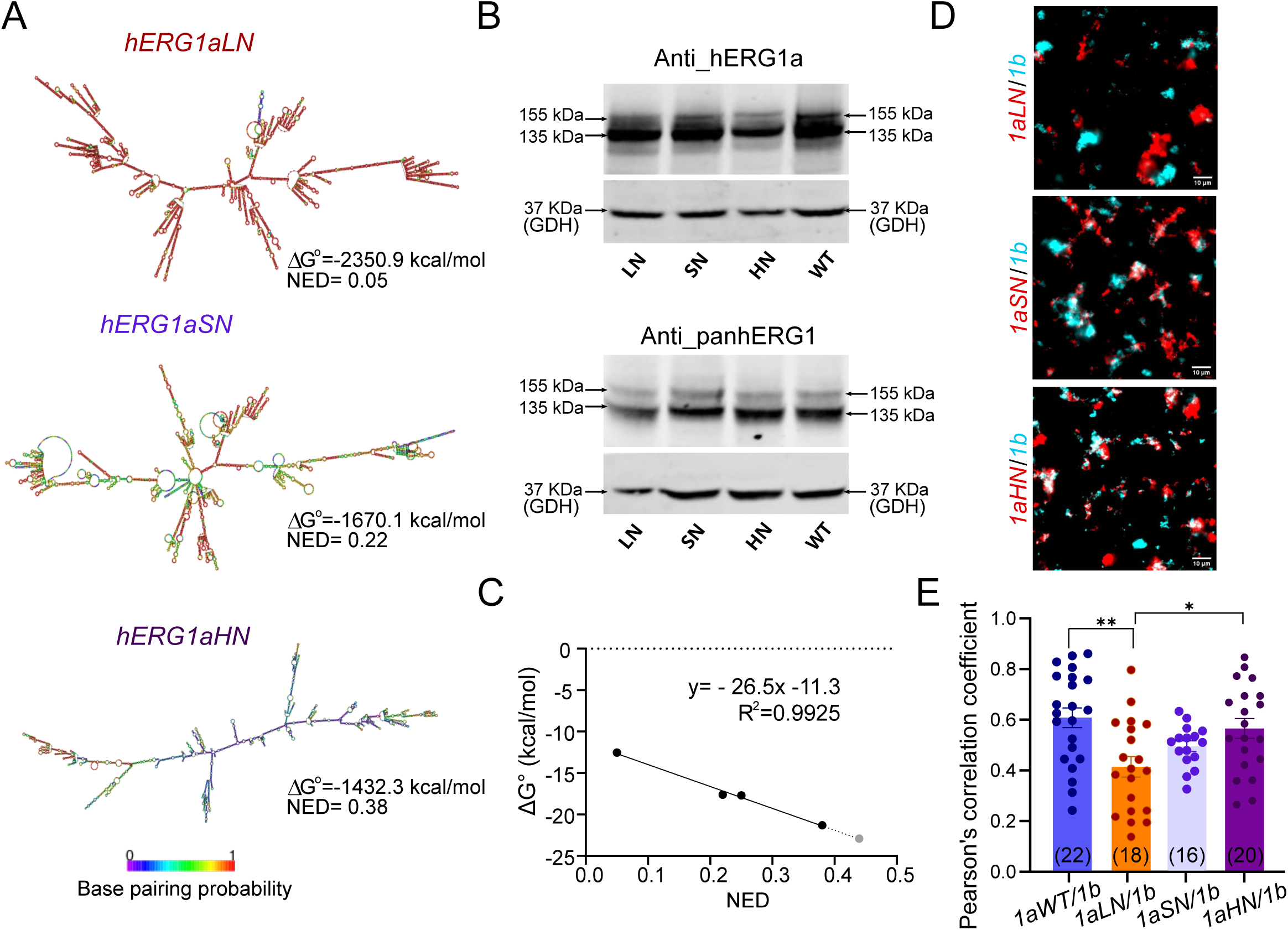
*hERG1a* structural diversity influences its association with *1b in vitro*. **A.** RNAfold-predicted secondary structure of mutant *hERG1a* mRNAs with 80% lower (top), similar (middle), and 1.5 times higher (bottom) NED than *hERG1aWT*. Color coding indicates base pairing probability. Predicted free energy of folding is shown for every mutant, highlighting the higher structural stability for mRNAs with lower NED. **B**. Western blot analyses with antibodies targeting hERG1a N-terminal and pan-hERG1 regions show wild-type and silent mutant *hERG1a* mRNAs encode appropriately sized mature (155 kDa) and immature (135 kDa) hERG1a proteins. Arrows and numbers indicate molecular weight of each band. GDH represents GAPDH used as a loading control. **C**. Negative linear relationship between the predicted *hERG1a/1b* free energy of binding and *hERG1a* mRNA NED. R^2^ and equation describing the obtained fit are shown. **D**. Representative images of *in vitro* heterotypic reconstitution of fluorescently labeled *hERG1b* and mutant *1a* mRNAs mixed in a 1:1 ratio to a final concentration of 40 nM at physiological K^+^ levels. Scale bars represent 10 µm. **E**. Comparison of the Pearson’s correlation coefficient of *hERG1a* and *1b* mRNA fluorescent signals in heterotypic mixes shown in **D**. Bars represent mean +^_^ SEM. **: p<0.01, *: p<0.05. Single points indicate independent experiments and numbers in parenthesis indicate replicates.

To examine whether *hERG1a* structural diversity influences its ability to directly interact with *1b*, we analyzed the relationship between *hERG1a* NED and the predicted free energy of binding (ΔG°) between the two transcripts (Fig 2C). This analysis revealed a negative linear relationship between both parameters, predicting that lowering *hERG1a* structural diversity would diminish its association with *1b*, while increased *hERG1a* NED would favor the mRNA interaction. To test this prediction, we reconstituted *hERG1b* with each mutant *hERG1a* mRNA and analyzed the relationship between Pearson’s correlation coefficient and *hERG1a* NED (Fig. 2D). As expected, the correlation coefficient was lower for *hERG1aLN/1b* mRNAs compared to the wild-type pair (0.41 v. 0.60), indicating that *hERG1b* associates less effectively with *1aLN* than with *1aWT* (Fig. 2E). The findings of this experiment strongly suggest that decreasing *hERG1a* structural diversity weakens its direct interaction with *1b*.

In agreement with our predictions, no significant changes were observed in the association of *hERG1b* with *1aSN,* as expected from its similar NED to *hERG1aWT* (Fig. 2E). However, *1aHN*, with its increased structural diversity predicted to enhance interactions with *hERG1b*, instead showed similar levels of association to *hERG1aWT* (Fig. 2E). This surprising finding can be explained by the possibility that, although a higher NED allows additional structural conformations that could favor *hERG1a* and *1b* interactions, the mutations introduced in *1aHN* disrupt sequences comprising specific sites mediating intermolecular interactions. Further work will be required to understand the impact of mutations in *hERG1aHN* on interactions with *hERG1b*.

### *hERG1a* structural diversity influences its association with *1b* in HeLa cells

To first determine whether the specificity of *hERG1a* and *1b* mRNA association is conserved in the cellular environment, we imaged live HeLa cells transfected with *in vitro*-transcribed, fluorescently labeled transcript pairs. Twenty-four hours after transfection, heterotypic condensates were observed (Fig. 3A, *SI Appendix,* Supplementary Video 1). These condensates were dynamic, undergoing multiple fusion and fission events characteristic of intracellular biomolecular condensates (47–49). mRNA colocalization was quantified for each frame of 40-second live-cell videos using a threshold distance of 300 nm between puncta centers. Colocalization is thus expressed as the percentage of the indicated mRNA complexes found in proximity to *hERG1b* complexes. This analysis revealed colocalization of 69.5% for *hERG1aWT* with *1b* and 15.6% for *PLN* with *1b*, reflecting a four-fold preference between *hERG1a* and *1b* mRNAs (Fig. 3B). This preference is greater in cells than *in vitro*, perhaps reflecting roles for other biomolecules in promoting *hERG1a* and *1b* association.

**Figure 3.**
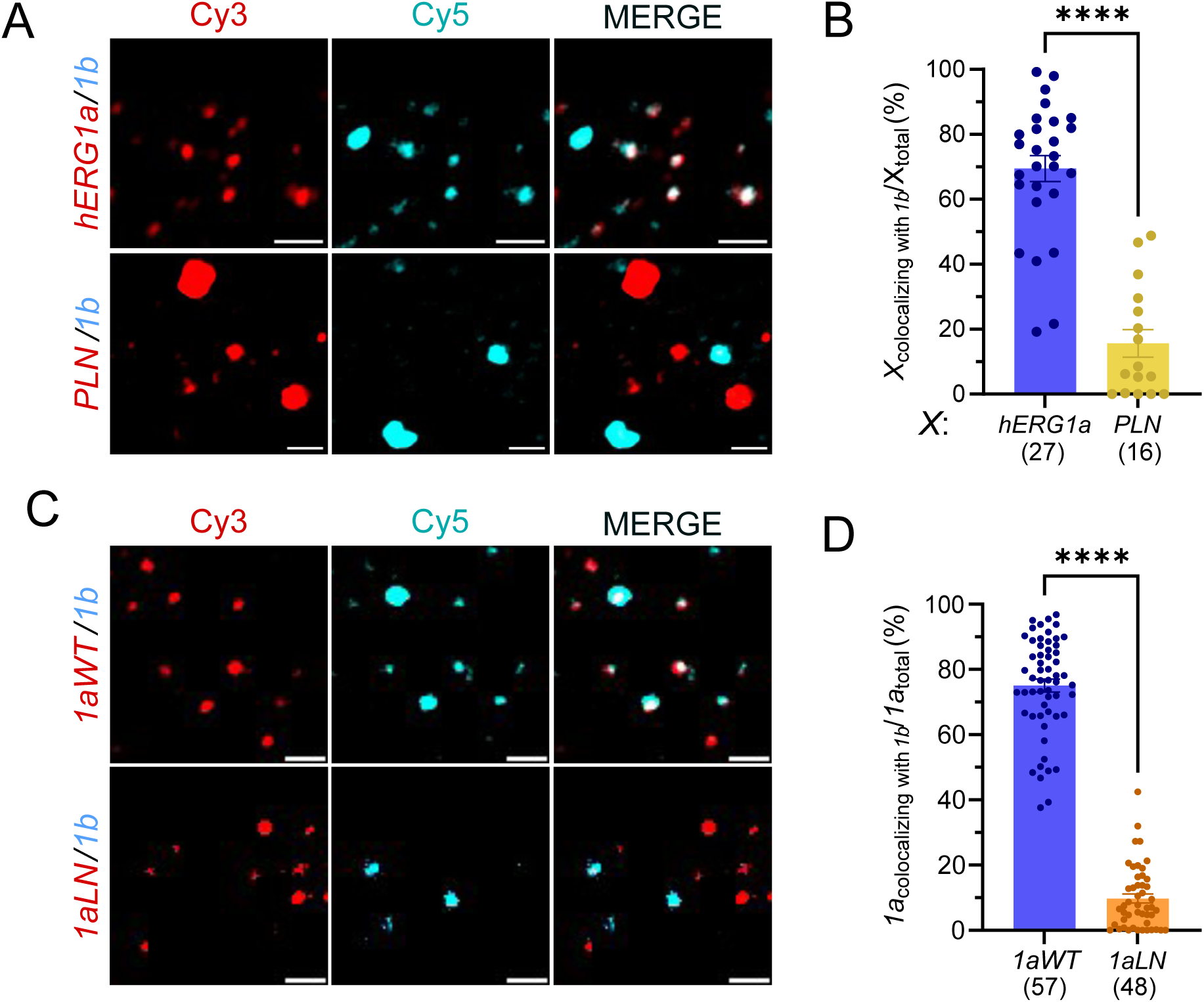
Specific *hERG1a* and *1b* mRNA interaction depends on RNA structural diversity in HeLa cells. **A.** Representative images of condensates formed 24 hours after cotransfecting *hERG1a/1b* or *PLN/1b* mRNA pairs in HeLa cells. **B**. Comparison of the percentage of colocalization of *hERG1a* or *PLN* mRNAs with *hERG1b* in live HeLa cells. **C**. Representative images of condensates formed 24 hours after cotransfecting *hERG1aWT/1b* or *hERG1aLN/1b* mRNA pairs in HeLa cells. **D**. Percentage of colocalization of *hERG1a* variants with *hERG1b* mRNA. Scale bars indicate 2 µm. Bars represent mean ^_^+ SEM. ****: p<0.0001. Single points indicate individual cells and numbers in parenthesis represent replicates.

To further test the hypothesis that mRNA physicochemical properties are responsible for *hERG1aWT* and *1b* overlap in HeLa cells, we determined the effect of reducing *hERG1a* structural diversity on intracellular transcript association (Fig. 3C). We analyzed the extent of the colocalization of *hERG1aWT or 1aLN* with *hERG1b* in 300-second live cell videos (Fig 3C, analysis for *1aSN* and *1aHN* in *SI Appendix*, Fig. S3). Cellular condensates exhibited markedly less overlap (10%) when *hERG1a* structural diversity was reduced, reflecting a 7-fold difference that is even more pronounced than the 1.5-fold difference observed *in vitro* (Fig. 3D). We conclude that reducing *hERG1a* structural diversity diminishes association with *1b* both *in vitro* and in the cellular environment, consistent with a role for direct mRNA interactions.

### mRNA association is important for concomitant translation of heteromeric subunits

If interaction of *hERG1a* and *1b* mRNAs is important for cotranslational heteromerization of their encoded subunits, then disrupting mRNA association by reducing structural diversity should interfere with heterotypic protein association. To test this hypothesis, we analyzed hERG1a translational complexes, defined by colocalization between mRNA and protein, in cells co-expressing *hERG1aWT* or *1aLN* with *hERG1b* (Fig. 4A, B). hERG1a and 1b subunits were C-terminally tagged with blue and green fluorescent proteins, respectively, without affecting the NED of the encoding transcripts (see Table 1 for NED comparison). Fluorescently labeled *hERG1a* and *1b* mRNAs encoding hERG1a-BFP and hERG1b-GFP, respectively, were cotransfected in HeLa cells and imaged 48 hours later. We found that 87% of *hERG1aWT* translational complexes were heterotypic, while 13% were homotypic, suggesting a greater efficiency of heteromeric vs. homomeric ion channel biogenesis. Treatment with puromycin, a translation inhibitor causing the dissociation of translating proteins from the ribosome (50), showed a reduction of 50% in *hERG1a* mRNA and protein colocalization (*SI Appendix*, Fig. S4). This reduction corresponds to that expected based on a previous study in which the concentration was titrated to achieve a 50% perturbation of translational complexes while maintaining cell viability (38, 51). This confirms that the identified complexes were actively translating condensates.

**Figure 4.**
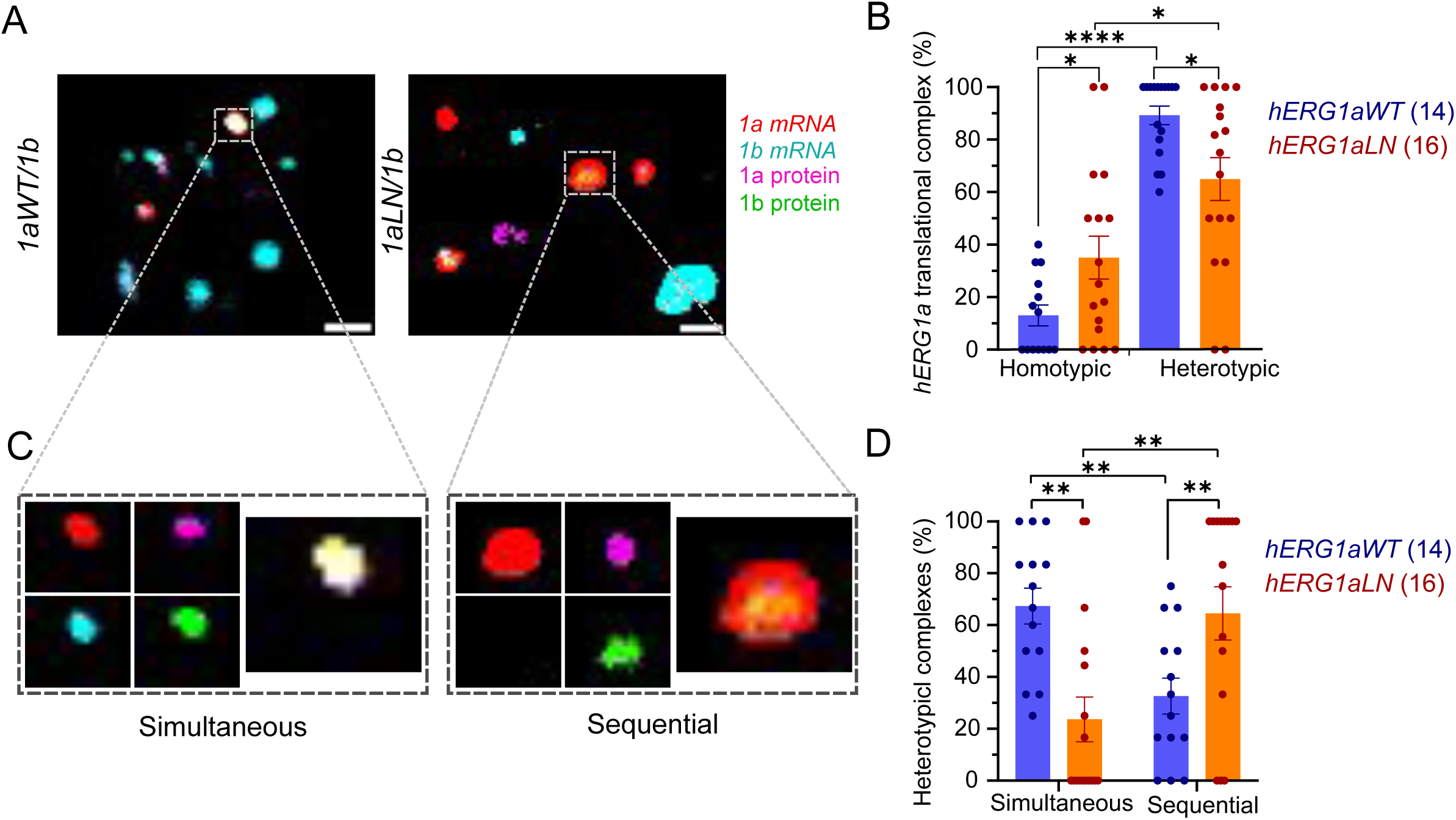
m RNA interaction influences hERG1a/1b co-translational association. **A.** Representative images of HeLa cells 48 hours after co-transfecting *hERG1b* with wild-type or mutant *hERG1a* mRNAs. *hERG1a* and *1b* mRNAs encode C-terminal fluorescently tagged hERG1a-BFP and hERG1b-GFP protein, respectively. Scale bar in images represent 2 µm. **B.** Classification of hERG1a translational complexes based on colocalization analysis. Homotypic complexes refer to colocalization between hERG1a mRNA and protein only, whereas heterotypic indicates colocalization of *hERG1a* mRNA with both proteins. **C.** Representative images of hERG1a/1b co-translational complexes in HeLA cells co-expressing *hERG1b* with *1aWT* or *1aLN*. **D.** Differential fractions of identiified co-transational complexes. Bars represent mean +^_^ SEM. ****: p<0.0001, **: p<0.01, *: p<0.05. Single points and numbers in parenthesis indicate individual cell replicates.

Substituting *hERG1aWT* with *hERG1aLN* to disrupt mRNA interactions reduced hERG1a and 1b subunit cotranslational heterotypic association by 30%. This reduction corresponded with an increase in homotypic hERG1a translational condensates (Fig. 4B). A quantitatively similar pattern emerged when analyzing *hERG1b* translational complexes (*SI Appendix*, Fig. S5). These results suggest that the fidelity of mRNA interactions is a critical factor underlying the biogenesis of ion channels in heterotypic complexes.

Upon further analysis, we were surprised to identify two distinct classes among cotranslational heterotypic complexes (Fig. 4C, D). One class is defined by the colocalization of *hERG1a* and *1b* mRNAs and their encoded proteins, suggesting the simultaneous translation of both mRNAs within a complex (Fig. 4C, *left*). A second class comprises only *hERG1a* mRNA colocalizing with both proteins, as if a nascent hERG1a subunit associates sequentially with a fully translated hERG1b protein (Fig 4C, *right*). A similar analysis performed in hiPSC-CMs cotransfected with fluorescently labeled *hERG1a* and *1b* mRNAs identified the same classes of translational complexes (*SI Appendix*, Fig. S6).

Quantification of WT heterotypic translational complexes reveals that 67% of nascent hERG1a subunits associate with *hERG1b* simultaneously, compared to 33% sequentially (Fig. 4D, violet bars). However, *hERG1aLN*, in addition to causing shift in biogenesis toward more homotypic complexes, yielded a marked reduction in simultaneous cotranslation with *1b* (to 27%), favoring instead sequentially cotranslated complexes (73%) (Fig 4D, orange bars). There were no differences in the fraction of mRNAs undergoing translation between *hERG1aWT* and *1aLN*, indicating that the observed changes were not due to reduced translation efficiency (*SI Appendix*, Fig. S7). These results suggest that the perturbation of mRNA association triggers a shift in the mode of coordinated cotranslational biogenesis.

## DISCUSSION

In this study, we observed using *in vitro* reconstitution that *hERG1a* and *1b* mRNAs interact directly to form heterotypic condensates, which were disrupted when the structural diversity, or NED, of *hERG1a* mRNA was reduced. In live cells, reduced NED diminished the formation of heterotypic, cotranslational condensates, promoting instead homotypic assemblies. Unexpectedly, the remaining heterotypic condensates showed fewer instances of simultaneously translated *hERG1a* and *1b* mRNAs. Instead, more condensates showed sequential cotranslation represented by both proteins but only *hERG1a* mRNA, as if the hERG1b protein had previously completed its synthesis. Our results highlight the relevance of mRNA structural diversity and association in steering the biogenesis pathway of heteromeric proteins.

Whereas changes in *in vitro* mRNA association were seen for reductions in NED, no significant differences were observed for increases in this parameter, in contrast to expectations from predictions of favorable free energy of binding of *hERG1aHN* and *1b*. A factor contributing to this outcome may be the inability to change the NED in equal proportions in either direction from WT. When altering *hERG1a* structural diversity, we allowed only synonymous mutations within the coding region that did not significantly affect the nucleotide content and fractional codon usage (*see Methods*). Given these constraints, a NED of 0.38 for *hERG1aHN* was the highest that could be achieved and may have been insufficient to increase mRNA association above *hERG1aWT*. However, as shown by the dashed gray line in Fig. 2C, a more favorable binding energy may have been realized were these limitations not in place.

Although increasing *hERG1a* NED may theoretically provide the potential for additional intermolecular interactions, the underlying mutations in *hERG1aHN* might have also disrupted specific sequence motifs critical for its association with *1b*, unexpectedly weakening the mRNA interaction. In cells, such changes in primary sequence may also disrupt interaction with other intracellular biomolecules such as RNA binding proteins or cytoplasmic small nucleolar RNAs (snoRNA), which may work in concert with direct interactions between the mRNA molecules (52, 53). These considerations highlight the limitations of considering only changes in the free energy of binding when altering the primary structure of RNA. Future studies will be required to identify specific interaction sites between the mRNAs and the role of other biomolecules during ion channel biogenesis.

Native mRNAs encoding functionally related proteins have previously been identified in pairs in ribonucleoprotein complexes that were smaller than the condensates observed in the current study (37, 38, 54). This difference may arise from the transfection of mRNA into the cells, which was designed to circumvent the involvement of mRNA cotranscription either *in vitro* or in cells and thus revealed that RNA physicochemical properties are sufficient to drive their association. Although further work is required to resolve this issue, we speculate that mRNA cotranscription and association with RNA binding proteins, as the transcripts exit the nucleus, might dictate the number of associating mRNAs and thus the size of the observed heterotypic cotranslational complexes.

A growing body of research suggests that protein complexes oligomerize through either simultaneous or sequential pathways, depending on the nature of their subunits and interactions (55, 56). Selective ribosome profiling in yeast has shown that soluble subunits engaging via N-terminal interactions often assemble during simultaneous translation, whereas those interacting through C-terminal regions favor sequential assembly (57). This observation aligns with our previous finding that hERG1a and 1b N-termini interact early in biogenesis (26), as well as our current observation of their concomitant translation within heterotypic complexes. Furthermore, a recent study combining ribosome profiling, structural analysis, and AI modeling proposed that protein complexes with highly intertwined subunits preferentially assemble via simultaneous cotranslation (56). Consistent with that report, our current study shows that hERG1 ion channels, characterized by extensive inter-subunit interfaces among transmembrane and intracellular domains (58), are mainly synthesized in simultaneous cotranslational complexes. Future studies using CRISPR-tagged subunits in cardiomyocytes will be essential to define the distribution of translation modes in more native cellular contexts.

Despite previous views that mRNAs in condensates such as stress granules and P-bodies are translationally repressed, recent reports challenge that conclusion (59). Studies in yeast have described granules in which specific mRNAs are translated during active growth conditions (60). Additionally, also in yeast, the association of fully translated and nascent proteins in condensates was proposed to facilitate the assembly of protein complexes (61). Our study reinforces the importance of condensates in translation, specifically in the biogenesis of heteromeric complexes.

Several studies have shown that synonymous mutations, once considered “silent,” can significantly impact translation efficiency, protein function, physiological roles, and pathogenesis (62). In the context of oligomeric assembly, for example, synonymous mutations in mRNAs encoding certain *E. coli* enzymes affect their translation rate and promote the formation of “entangled” domains at dimerization interfaces as modeled by molecular dynamics simulations (63). Similarly, synonymous mutations in the mRNA encoding chloramphenicol acetyltransferase, an enzyme essential for *E. coli* growth in the presence of chloramphenicol, perturb cotranslational folding and make the encoded protein susceptible to degradation, impairing bacteria growth (64). In human neurons, a synonymous single nucleotide polymorphism (C957T) in *DRD2* mRNA alters its secondary structure, stability, and translation efficiency, reducing expression of the D2 dopamine receptor (65). Synonymous mutations can also contribute to cancer, as seen in *CDKN2B* and *BRCA2* mRNAs, where disrupted post-transcriptional mRNA modification enhances tumorigenesis (66). Indeed, in hERG1a, synonymous mutations designed to increase expression paradoxically reduced mRNA stability, lowering protein levels and altering channel function (67). With the increasing body of literature highlighting the functional and pathophysiological impact of synonymous mutations, we anticipate greater focus in future clinical studies to link synonymous mutations to long QT syndrome and other diseases.

## METHODS

### Generation of mutant mRNAs and thermodynamic analyses

Structural mutants of *hERG1a* mRNA were designed by creating a genetic algorithm that uses secondary structure and RNA-RNA interaction prediction tools from the ViennaRNA package (68). This algorithm iteratively creates generations of synonymously mutated sequences until an optimal one is found. Every given generation results in a library of 128 sequences, each of which has between 5 and 50 codons swapped at random locations within the coding sequence. Only sequences whose nucleotide content and fractional codon usage vary by less than 1% and 10%, respectively, compared to *hERG1aWT*, remain in the population. Mutants with different structural content were chosen based on their predicted minimum free energy (MFE) in RNAfold using the full mRNA sequence, including the 3’UTR (68). The choice of the optimal mutant sequence is subject to the Metropolis criterion. Python scripts ‘evolvehERG1a_struc_match_codon_CLUSTER.py’ and ‘evolvehERG1a_unstruc_match_codon_CLUSTER.py’ implementing the parallelized genetic algorithm were run on single nodes using 124 CPUs each on the Longleaf computer cluster at the University of North Carolina at Chapel Hill. RNAup from the ViennaRNA package was used to determine the free energy of binding between the resultant mutant *hERG1a* mRNAs and *hERG1b,* considering the free energy for the dimerization and conformational changes required for each mRNA (68). Mutant *hERG1a* mRNAs were synthesized and cloned in pcDNA 3.1 using Genescript service.

### *In vitro* transcription and labeling of mRNAs

*PLN, hERG1a* variants, and *1b* coding sequences and 3’UTRs were cloned in a pcDNA3.1 backbone. All pCDNA3.1 plasmids contained a T7 promoter for mRNA *in vitro* transcription. Plasmids containing *hERG1b* and *1a* variants (WT, SN, and HN) were linearized with EcoRI restriction enzyme at the 3’end, whereas those containing *hERG1aLN* and *PLN* were linearized with EcorV and XhoI, respectively. Linearized plasmids were transcribed using mMESAGE mMACHINE™ T7 Transcription Kit (Invitrogen, AM1344) and fluorescently labeled with Cy3 or Cy5 with Label IT nucleic acid labeling kit (Mirus, MIR3600 and MIR3700). Agarose gels were run to check for mRNA integrity after every step. mRNA concentration was quantified using a nanodrop (Thermo Fischer).

### *In vitro* reconstitution, imaging, and analysis

*In vitro* transcribed and fluorescently labeled mRNAs were mixed in a 1:1 ratio to the final concentration indicated in each set of experiments. Mixed mRNAs were co-incubated for 3 hours at 37⁰C in buffers containing fixed 10 mM MgCl_2_ and varying concentrations of spermine and KCl. Spermine (Sigma) concentration ranged from 0 to 10 mM while KCl ranged from 0 to 200 mM. Reconstituted mixes were imaged in Zeiss Axiovert 200 inverted microscope (Plan-apochromatic, air, 20X, 0.17 NA). Images were analyzed via Image J Fiji (https://imagej.net), and Pearson’s correlation coefficient was calculated with the plugin JACOP (https://imagej.net/ij/plugins/track/jacop2.html).

### Cell culture and transfection

HEK-293 and HeLa cells (ATCC) were cultured using DMEM culture media (ThermoFisher, 11995065) supplemented with 10% FBS (ATCC, SCRR-30-2020). For western blot analysis, HEK-293 cells were plated in 35 mm plates, and transient transfections were performed using 2.5 μL/mL Lipofectamine 2000 (Thermofisher) with 1 μg of plasmid. For live cell imaging, HeLa cells were cultured in 35mm glass bottom dishes with #1.5 cover glass (Cellvis, D35-14-1.5-N) and transfected with a 1:1 ratio of mixed *in vitro* transcribed and fluorescently labeled mRNAs to a final concentration of 500 ng/mL. mRNA transfection was performed using a Trans-IT-mRNA transfection kit (Mirus MIR 2225). Human iCell iPSC-CMs (CDI-FujiFilm) were thawed, plated, and cultured with CDI media according to the manufacturer’s specifications. After 10 to 14 days in culture, cells were transfected using Trans-IT-mRNA transfection kit and imaged 24 hours later. For experiments testing translation inhibition, HeLa cells were treated with 100 µM of puromycin 1or 18-hours before imaging as indicated.

### Western Blot

HEK-293 cells were collected 48 hours post-transfection. Cell lysates were mixed with 4X loading buffer to a 1X final concentration and 20 nM dithiothreitol (DTT) and denatured at 50 °C for 10 minutes before loading on a 4-20% precast polyacrylamide gel. The gel was run at 100 V for approximately 120 minutes and transferred to PVDF membrane using standard protocols. Blots were blocked in 5% milk in TBS-T and probed overnight with hERG1a (Cell Signaling Technology, hERG1a (D1Y2J) Rabbit mAb #12889) and pan-hERG (ENZO Life Sciences, ALX-215-049-R100 HERG1 (CT) (pan) polyclonal antibody) antibodies. The membranes were washed three times for 5 minutes with TBS-T and incubated for 1 hour with appropriate secondary antibodies (1:10,000). Blots were washed twice for 5 minutes and imaged using a ChemiDoc MP Imaging System (BioRad).

### Live cell imaging and analysis

For live cell imaging, HeLa cells were transfected with fluorescently labeled mRNAs. To determine intracellular mRNA association, cells were imaged 24 hours post-transfection using a Nikon Ti2 spinning disk confocal microscope with a 60x (1.4 NA) oil immersion objective and Hamamatsu ORCA-Flash4.0 sCMOS camera. To analyze hERG1 translational complexes, cells were imaged using a Nikon AXR inverted point scanning confocal microscope with a 60X PLAN APO lambda D (1.42 NA) oil immersion objective with a high sensitivity GaASP detector. Microscopes were connected to a Tokei Hit enclosure for live cell imaging, where cells were kept at 37C °C and 5% CO2. Nikon Elements software was used for image acquisition.

Puncta determination and colocalization analysis were performed using the Imaris (Oxford instruments) spots model. Puncta were detected throughout the field of view of the microscopy image, and the colocalization analysis was performed for every cell individually. For the analysis of mRNA association, 300 nm between the puncta centers was used as a threshold, whereas for the classification of translational complexes, the threshold was 400 nm. Distances between the puncta centers were calculated using the Vantage plugin of Imaris (Oxford instruments). To ensure unbiased analysis, blinded technicians performed puncta determination and colocalization analyses using configuration files containing thresholding parameters across images.

All the quantifications were conducted on raw images. The sample images provided in the figures were deconvolved using Huygens professional software (Scientific Volume Imaging) and the Classic Maximum Likelihood Estimation (CMLE) algorithm with default settings.

### Statistical analysis

Statistical analyses were performed using Microsoft Excel and GraphPad Prism 9. To compare different conditions, we used the Student’s T-test or one-way ANOVA. All the data are shown in mean ± S.E.M.

## ACKNOWLEDGEMENTS

We thank Zachary T. Campbell and members of the Robertson lab, especially Sudharsan Kannan, and Taylor Voelker for critical discussion. The work was supported by NIH grants NHLBI 1R01HL131403-01A1 (G.A.R.) and K99HL169909 (L.F.A.). I.S. thanks the Max Planck Society and the Alexander von Humboldt Foundation for financial support.

## Author contributions

L.F.A. and G.A.R designed research; L.F.A, A.S.H, and F.L. performed experiments; L.F.A, I.S, and A.S.G. designed mRNA constructs; L.F.A. and L.C. analyzed data; and L.F.A. and G.A.R. wrote the paper.

## Classification

Biological Sciences

---

**Supplementary Figure 1.**
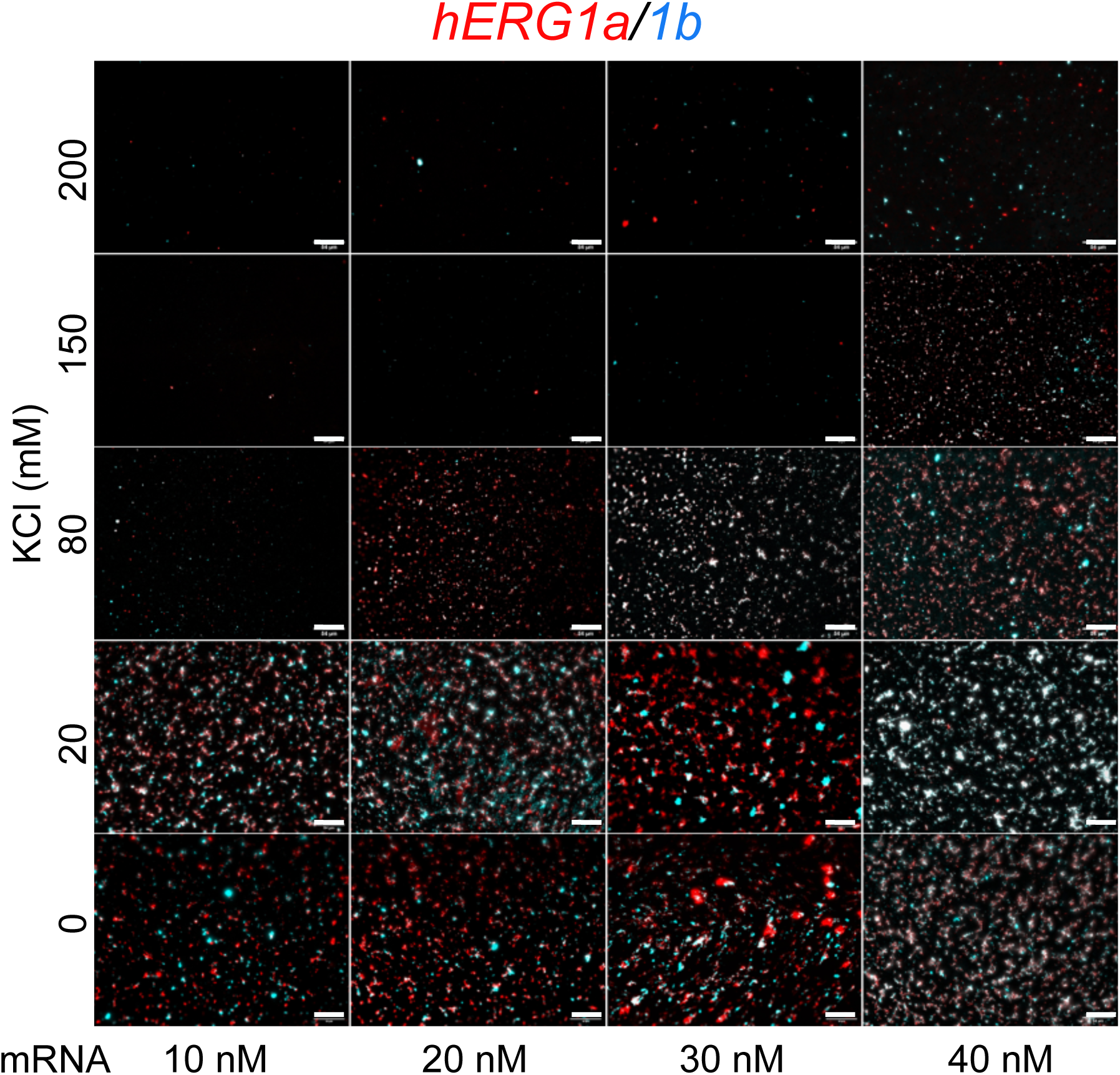
Representative images of *in vitro* reconstituted *hERG1a* and *1b* mRNAs. Different concentrations of *in vitro* transcribed and fluorescently labeled *hERG1a* and *1b* mRNAs were reconstituted in a 1:1 ratio in distinct KCl concentrations. Scale bars represent 50 μm. Increasing the total mRNA concentration visually augmented the area convered by condensates at fixed KCl concentration, as expected due to higher crowding effect. Interestingly, changes in ionic strength via increased KCl concentration visibly diminished the area covered by condensates, suggesting that changes in salt levels affect the mRNA intermolecular associations. These results suggest that the association between *hERG1a* and *1b* mRNAs in microscopic complexes is modulated by the microenvironment as previously reported for other biomolecular condensates reconstituted *in vitro* (39, 40).

**Supplementary Figure 2.**
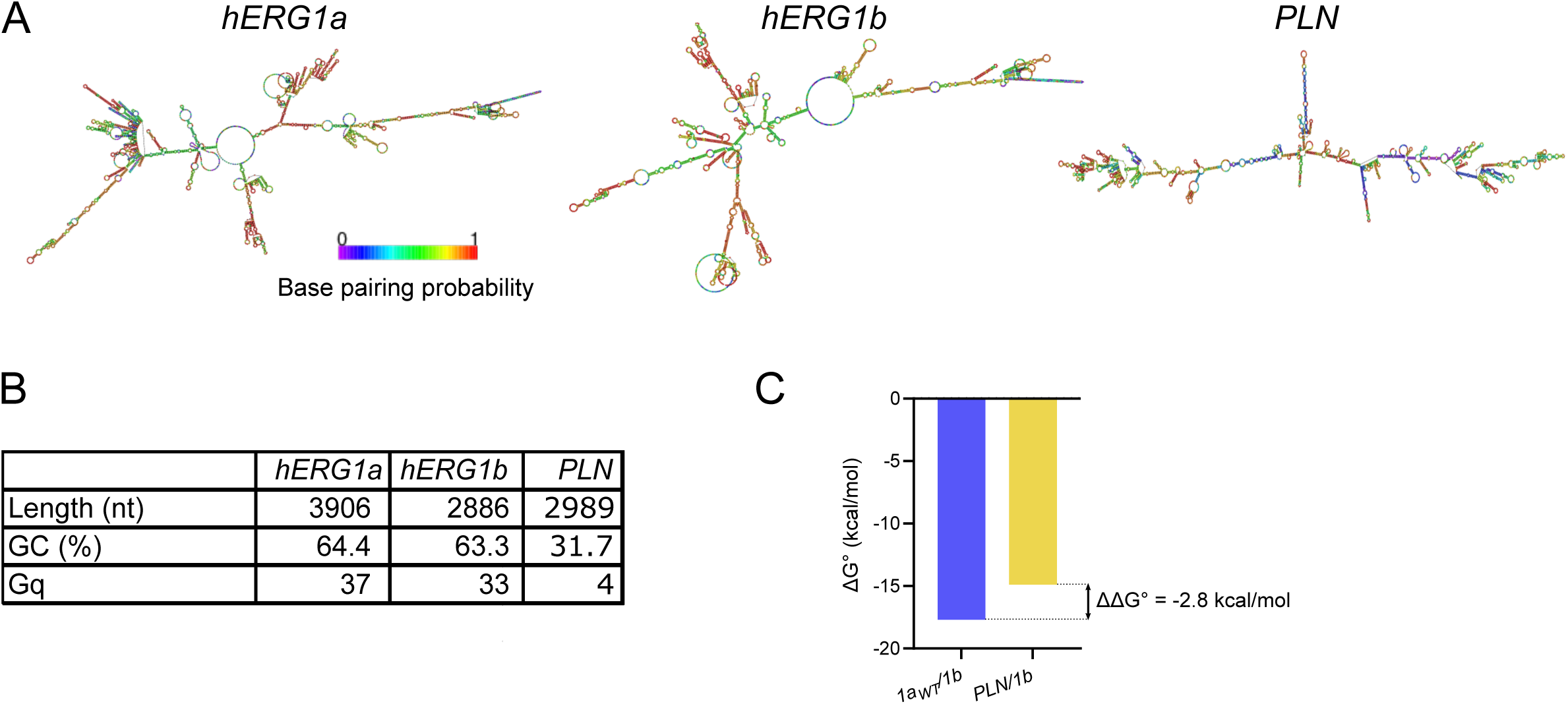
Computational analyses predict thermodynamically favorable direct interactions between *hERG1a* and *1b* mRNAs. **A.** RNAfold-predicted secondary structure of *hERG1a*, *1b*, and *PLN* mRNAs. Color coding indicates base pairing probability. For more details see *Methods* **B.** Table summarizing sequence and structure analyses of *hERG1a*, *1b*, and *PLN* mRNAs. **C**. Comparison of the predicted free energy of binding of *hERG1b* with *hERG1a* or *PLN* mRNAs. The negative ΔΔG° suggests association between *hERG1a* and *1b* mRNAs is thermodynamically favored compared to the *hERG1b/PLN* mRNA pair.

**Supplementary Figure 3.**
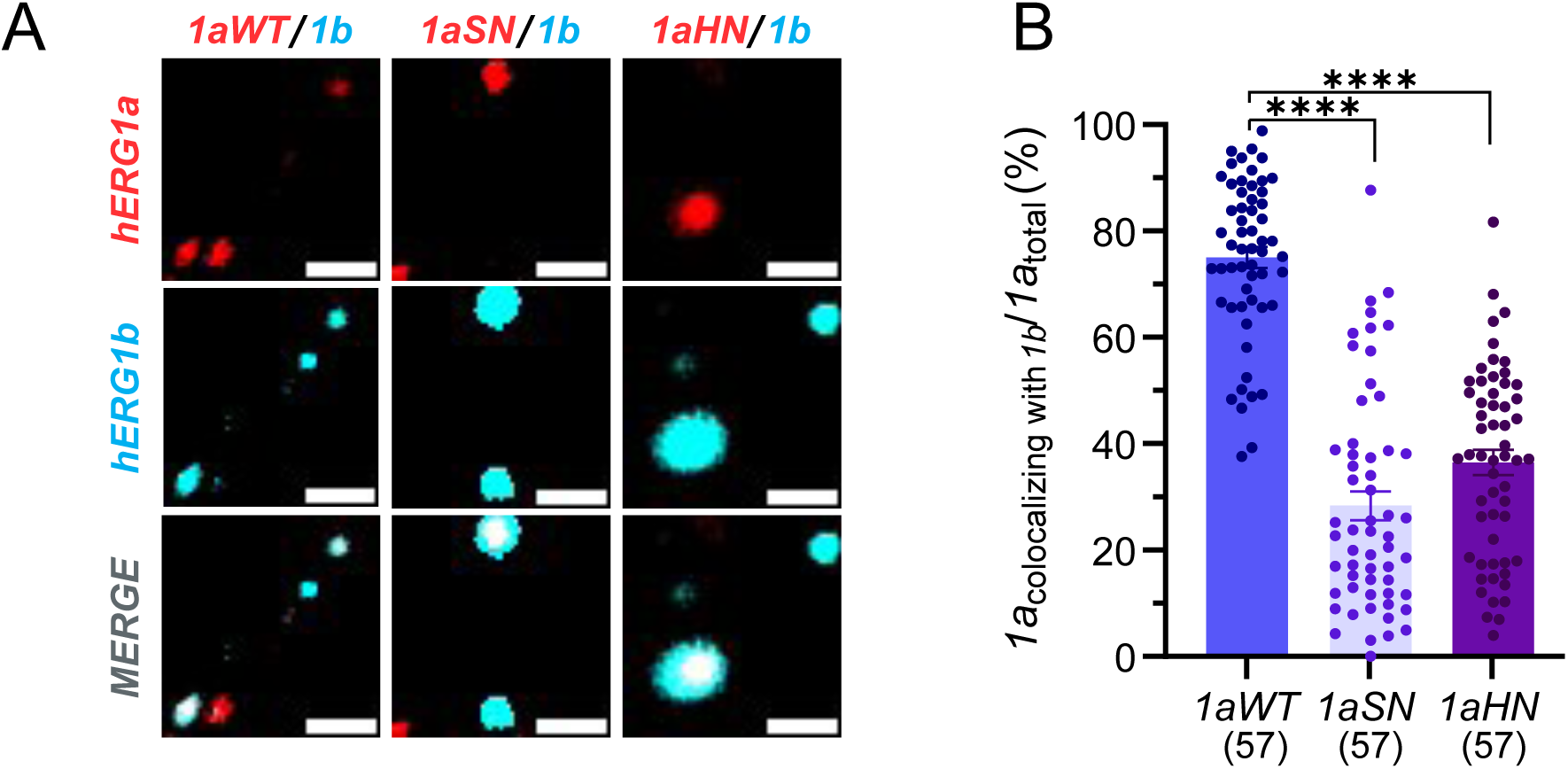
*hERG1a* sequence and structure influence its association with *1b* in HeLa cells. **A**. Representative HeLa cells imaged 24 hours post-transfection of independently *in vitro* transcribed and fluorescently labeled *hERG1aWT/1b*, *1aSN/1b,* and *1aHN/1b* mRNA pairs. mRNAs formed intracellular condensates with different heterotypic compositions. Scale bars represent 2 µm. **B.** Comparison of the colocalization percentage of cotransfected mRNAs based on a threshold distance of 300 nm between puncta centers. In contrast to *in vitro* data, these results suggest that changes in mRNA sequence is sufficient to disrupt *hERG1a* and *1b* mRNA association in the cellular environment. Bars represent mean ± SEM. ****: p<0.0001. Single points indicate individual cells and numbers in parenthesis represent replicates.

**Supplementary Figure 4.**
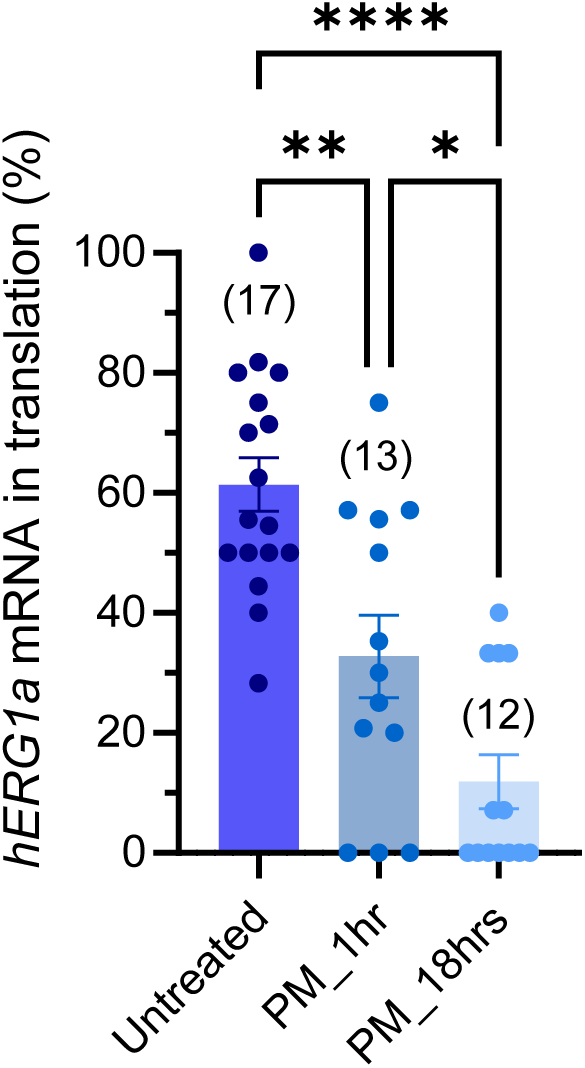
hERG1a mRNA and protein associate in translationally active condensates in HeLa cells. Comparison of the colocalization between hERG1a mRNA and protein in mock or puromycin-treated HeLa cells. Treatment with 100 μM puromicin decreased the colocalization between *hERG1a* mRNA and the encoded protein, suggesting that the observed intracellular condensates are translationally active. Bars represent mean ± SEM. *: p<0.5, **: p<0.01. ****: p<0.0001. Single points and numbers in parenthesis indicate individual cells.

**Supplementary Figure 5.**
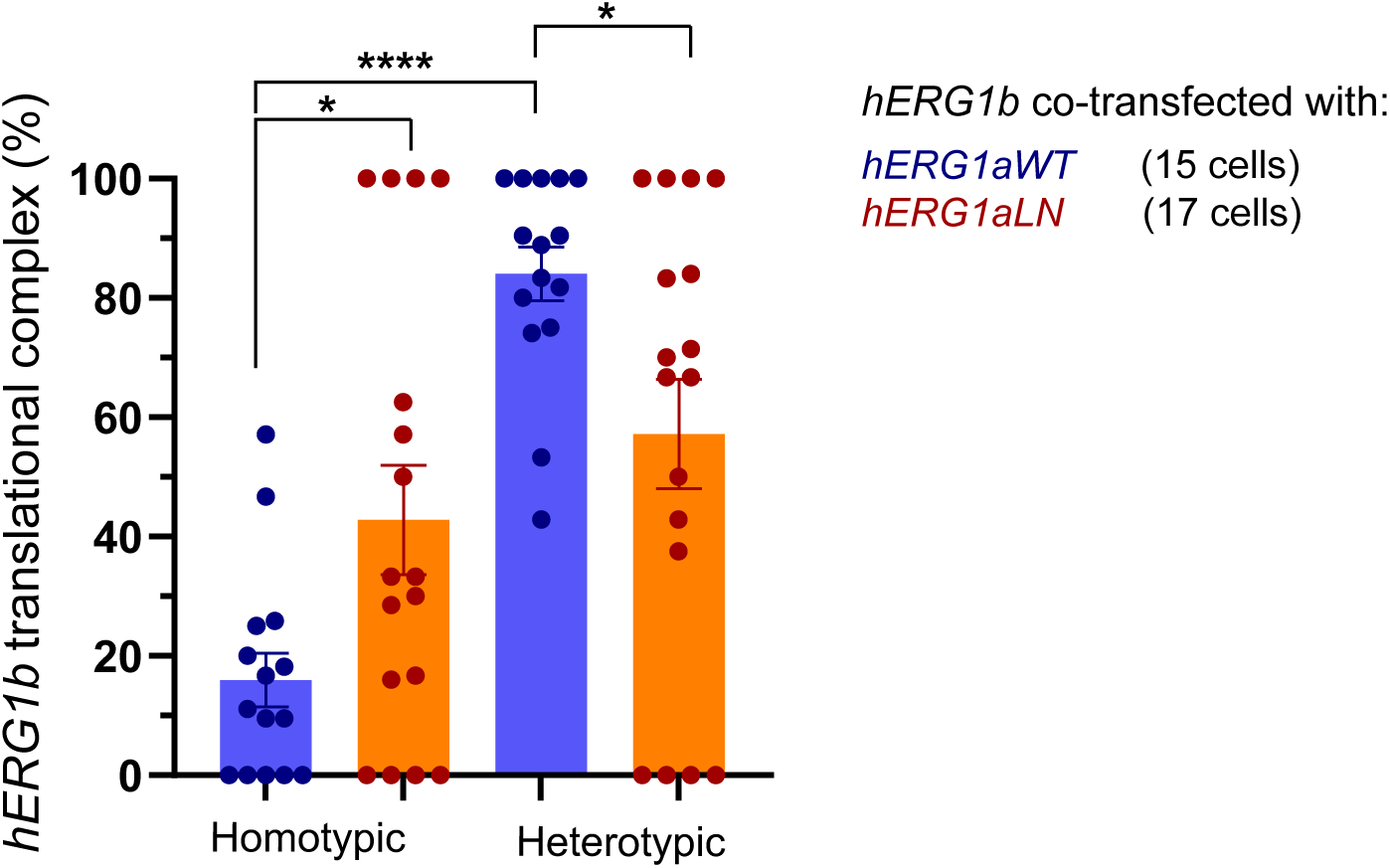
m RNA interaction is important for cotranslational heteromeric association. Classification of hERG1b translational complexes based on colocalization analysis. Homotypic complexes refer to colocalization between hERG1b mRNA and protein only, whereas heterotypic indicates colocalization of *hERG1b* mRNA with hERG1a and 1b proteins. Mutant *hERG1a* mRNA showing decreased association with *1b in vitro* and in cells increased the translation of hERG1b subunits in homotypic complexes, with consequent decrease of its cotranslational association with hERG1a. Bars represent mean +_ SEM. ****: p<0.0001,*: p<0.05. Single points indicate individual cells.

**Supplementary Figure 6.**
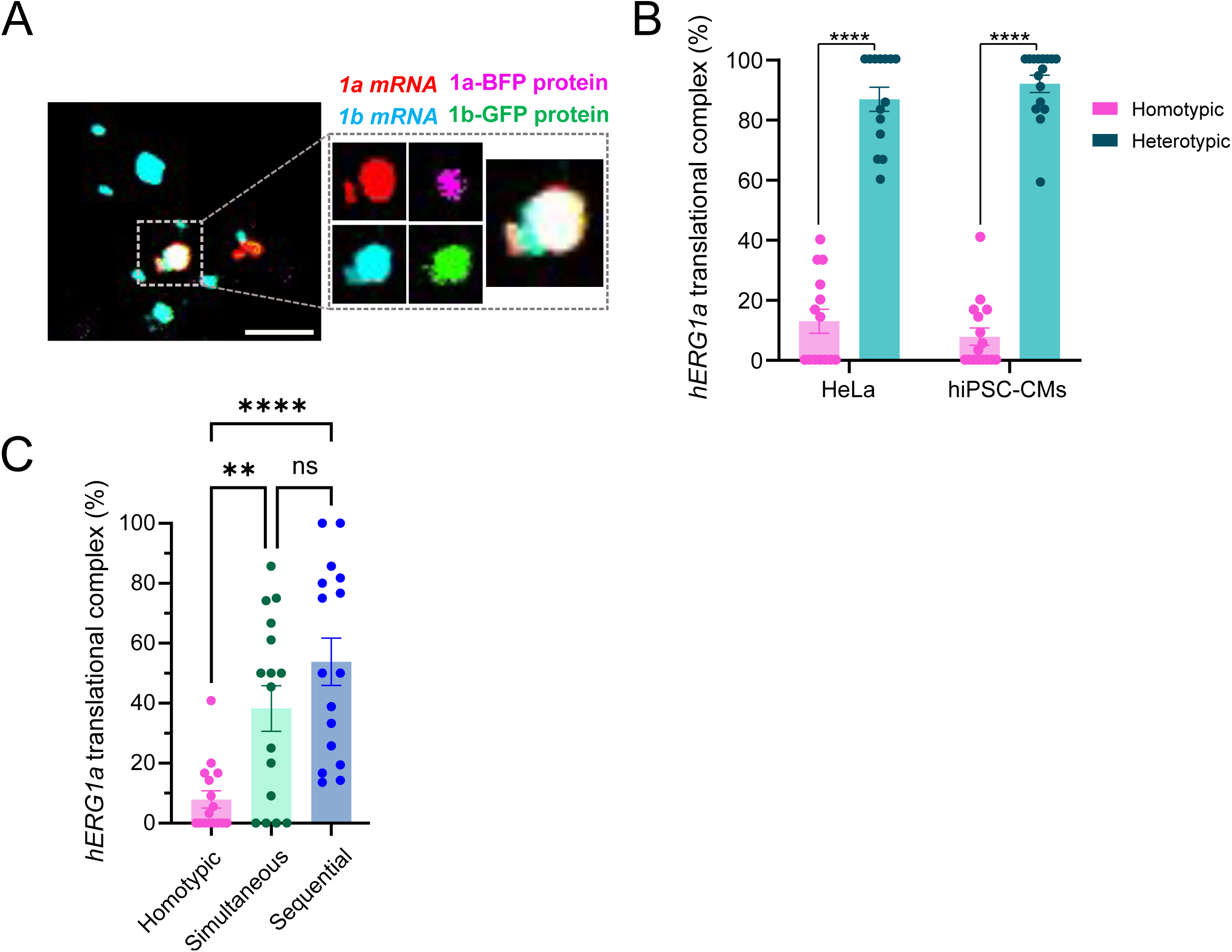
hERG1a/1b biogenesis occurs via diverse co translational pathways in hiPSC-CMs. **A.** Representative image of hiPSC-CMs 48 hours post cotransfection of fluorescently labeled *hERG1a* and *1b* mRNAs encoding hERG1a-BFP and hERG1b-GFP, respectively . Scale bar in images represent 5 µm. **B.** Comparison between homo- and heterotypic *hERG1a* translational complexes in HeLa and hiPSC -CMs. Similarly to translational complexes reconstituted in HeLa cells, *hERG1a* mRNA is predominantly translated in heterotypic complexes where hERG1b protein is also present. **C.** Classification of *hERG1a* translational complexes. Classification of cotranslational complexes in cardiomyocytes should be carefully interpreted as in these experimental conditions native hERG1a and 1b mRNAs and proteins are present and unlabled. Bars represent mean ± SEM. **: p<0.01, ****: p<0.0001. Single points indicate individual cells. A total of 16 hiPSC-CMs from three biological replicates were analyzed.

**Supplementary Figure 7.**
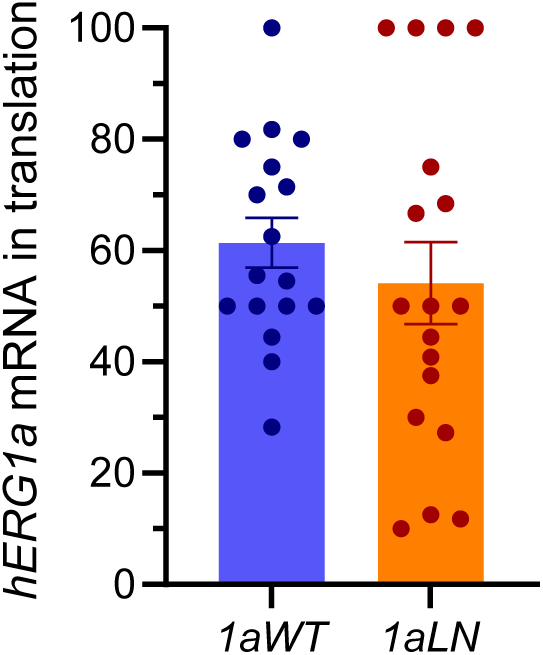
Synonymous mutations decreasing *hERG1a* NED do not reduce its translation efficiency. Comparison of the percentage of *hERG1a* mRNA undergoing translation based on colocalization analyses (hERG1a mRNA and protein) shows similar average values between wild-type and mutant *hERG1a* mRNAs. Individual points represent individual cells.

