## Supplementary File 1 for "mRNA interactions promote cotranslational association of heteromeric membrane proteins"

Supplementary File 1. Sequences corresponding to synonymous mutated mRNAs.

#### hERG1aLN

ATGCCGGTGCGGAGGGGCCACGTCGCCCCGCAGAACACCTTCCTGGACACCATCATCCGGAAGTTCGAAGGTCAGTCCAGGA
AGTTCATTATCGCCAACGCTAGGGTGGAGAACTGCGCCGTCATCTACTGTAACGACGGCTTCTGCGAGCTGTGCGGGTACTCGC
GGGCTGAGGTGATGCAGCGGCCCTGCACCTGCGACTTCCTGCACGGCCCCCGCACCCAGCGGCGGGCTGCCGCCCAGATAG
CCCAGGCCCTGCTGGGCGCCGAGGAGCGTAAGGTCGAGATCGCGTTCTACCGGAAGGACGGCTCATGCTTCCTGTGTCTGGT
GGACGTGGTCCCCGTGAAGAACGAGGACGGGGCTGTGATCATGTTCATCTTGAATTTCGAGGTCGTGATGGAGAAGGACATGG
TCGGCAGCCCCGCTCACGACACCAACCACCGCGGCCCGCCGACCAGCTGGCTGGCCCCCGGCCGGGCGAAGACCTTCCGCC
TGAAGCTGCCGGCGCTGCTGGCCCTCACCGCCCGAGAGAGCTCGGTGCGGTCCGGCGGCGCCGGCGGCGCCGGGGCGCCA
GGGGCCGTGGTGGTGGACGTGGACCTTACGCCCGCGGCGCCCAGCAGCGAGAGCCTCGCTCTCGACGAGGTCACCGCCATG
GACAACCACGTGGCGGGCCTGGGCCCGGCGGAGGAGCGGAGGGCTCTCGTGGGGCCCGGCAGTCCGCCCAGGAGTGCCCC
GGGGCAGCTGCCCTCGCCCCGGGCGCACTCGCTGAACCCGGACGCGTCGGGCTCGAGCTGCAGCCTCGCCCGGACGCGCA
GCCGAGAGAGCTGCGCGTCGGTGAGGCGCGCCTCGAGCGCCGACGACATCGAGGCGATGAGGGCGGGGGTGCTCCCACCC
CCCCCTCGCCACGCCTCGACGGGCGCCATGCACCCCCTCCGCTCCGGCTTGCTGAACAGCACGTCGGACTCGGACCTGGTGC
GCTACCGCACCATCTCGAAGATCCCTCAGATCACACTGAACTTCGTGGATCTGAAGGGCGACCCCTTCCTGGCTTCGCCCACCT
CCGACCGGGAGATCATCGCTCCCAAGATAAAGGAGCGCACCCACAACGTCACCGAGAAGGTGACTCAGGTGCTGAGCCTCGG
CGCCGACGTGCTGCCAGAGTACAAGCTGCAGGCGCCGCGCATCCACCGATGGACCATCCTCCATTACAGCCCGTTCAAGGCCG
TGTGGGACTGGCTCATTCTTCTGCTGGTCATCTACACGGCCGTGTTCACCCCCTACTCGGCTGCCTTCCTTCTCAAGGAGACCG
AGGAGGGGCCGCCTGCCACCGAGTGTGGCTACGCCTGCCAGCCGCTAGCCGTCGTGGATCTGATCGTGGACATCATGTTCATT
GTGGACATCCTGATCAATTTCAGGACCACCTATGTCAACGCCAACGAGGAGGTGGTGAGCCACCCCGGCCGCATCGCGGTCCA
CTACTTCAAGGGGTGGTTCCTCATCGACATGGTGGCAGCAATCCCCTTCGACCTGCTGATCTTCGGCAGCGGGTCGGAGGAGC
TCATCGGGTTGCTGAAAACCGCCCGCCTGCTGCGCCTAGTGCGTGTCGCGCGAAAGCTGGATCGCTACAGCGAATACGGCGCT
GCTGTGCTGTTTTTGCTGATGTGCACCTTCGCGCTGATCGCGCACTGGCTGGCTTGCATCTGGTACGCCATAGGTAACATGGAG
CAGCCGCATATGGACTCCCGCATCGGCTGGCTCCACAACCTAGGCGACCAGATCGGCAAGCCATACAACAGCAGCGGGCTGGG
CGGTCCCTCGATCAAGGACAAGTACGTGACGGCGCTGTACTTCACCTTCAGCTCGCTCACCTCCGTGGGCTTTGGCAACGTCT
CTCCGAACACCAACTCGGAGAAGATCTTCAGCATCTGCGTGATGCTGATCGGGTCGCTCATGTATGCCAGCATCTTTGGCAACG
TGAGCGCCATAATCCAGCGGTTGTACTCCGGCACCGCTCGGTATCATACACAGATGCTGAGGGTGCGGGAGTTCATCCGCTTTC
ACCAGATCCCGAACCCGCTCCGGCAGCGGCTGGAGGAGTACTTCCAGCACGCGTGGAGCTACACGAACGGCATCGACATGAAT
GCAGTGCTGAAGGGGTTCCCTGAGTGCCTGCAGGCAGACATCTGCCTGCACCTCAACCGCTCACTCCTACAGCACTGCAAGCC
GTTCCGCGGCGCCACCAAGGGCTGTCTGCGGGCGCTGGCCATGAAGTTCAAGACCACCCACGCCCCACCCGGGGATACCTTG
GTCCACGCCGGCGACCTGCTGACAGCCCTTTACTTCATTAGCCGGGGCAGCATCGAGATCCTGCGGGGCGACGTGGTCGTGG
CCATCCTGGGGAAGAACGACATCTTCGGGGAGCCCCTGAACCTGTACGCACGCCCCGGCAAGAGCAACGGGGACGTGCGTGC
ACTTACGTATTGTGACCTCCACAAGATCCACCGCGACGATCTCCTGGAGGTCCTCGATATGTACCCCGAATTCTCGGACCACTTC
TGGTCCAGCCTGGAGATCACGTTCAACCTCCGGGACACGAACATGATCCCAGGCAGCCCGGGCTCGACCGAGCTCGAGGGCG
GGTTCAGCCGCCAACGGAAGCGCAAGCTCTCGTTCCGGCGGCGCACGGACAAGGACACGGAGCAGCCGGGGGAGGTGTCTG
CGCTGGGGCCGGGCCGGGCGGGTGCAGGGCCGAGTAGCAGGGGGCGGCCCGGGGGGCCGTGGGGGAGAGCCCGAGCT
CGGGGCCCTCGTCCCCTGAGTCATCAGAGGACGAGGGCCCCGGCCGGAGCTCCTCCCCCCTGCGGCTGGTTCCCTTCTCCA
GCCCGCGGCCCCCCGGGGAGCCCCCCGGGGGGGAGCCCCTGATGGAGGATTGCGAGAAGTCTAGTGACACCTGCAATCCTC
TGTCGGGGGCTTTCTCAGGAGTGAGTAACATCTTCTCCTTCTGGGGGGACTCCCGGGGGCGGCAGTACCAGGAGCTCCCTCG
GTGCCCTGCCCCCACCCCCAGCCTGTTGAACATCCCGCTCTCCAGCCCGGGGCGCCGGCCCCGCGGCGATGTGGAGAGCAG
GCTGGACGCTCTCCAGCGCCAGCTCAACAGGCTGGAAACCCGCCTGTCGGCAGACATGGCGACGGTGCTGCAGCTGCTGCAG
CGCCAGATGACTCTGGTCCCCCCTGCCTACTCGGCCGTGACCACCCCCGGGCCCGGCCCCACCAGCACCTCCCCCCTGCTGC
CCGTGTCCCCGCTGCCGACCCTAACCCTCGACAGCTTGTCGCAGGTCAGCCAATTCATGGCCTGCGAGGAGCTGCCCCCAGG
GGCCCCCGAACTTCCCCAGGAAGGCCCGACTCGCCGCCTCTCCCTACCGGGCCAGCTGGGCGCCCTCACCAGCCAGCCCCT
GCATCGCCACGGCTCCGACCCGGGCAGCTAGTGGGGCTGCCCAGTGTGGACACGTGGCTCACCCAGGGATCAAGGCGCTGC
TGGGCCGCTCCCCTTGGAGGCCCTGCTCAGGAGGCCCTGACCGTGGAAGGGGAGAGGAACTCGAAAGCACAGCTCCTCCCC
CAGCCCTTGGGACCATCTTCTCCTGCAGTCCCCTGGGCCCCAGTGAGAGGGGCAGGGGCAGGGCCGGCAGTAGGTGGGGCC
TGTGGTCCCCCCACTGCCCTGAGGGCATTAGCTGGTCTAACTGCCCGGAGGCACCCGGCCCTGGGCCTTAGGCACCTCAAGG
ACTTTTCTGCTATTTACTGCTCTTATTGTTAAGGATAATAATTAAGGATCATATGAATAATTAATGAAGATGCTGATGACTATGAATAAT
AAATAATTATCCTGAGGAGACTCCAAAAAAAAAAAAAAAAAA

### hERG1aSN

ATGCCGGTGCGGAGGGGCCACGTCGCGCCGCAGAACACCTTCCTGGACACCATCATCCGCAAGTTTGAGGGCCAGAGCCGTA
AATTTATCATCGCCAACGCTCGGGTAGAAAACTGTGCAGTAATTTATTGTAACGACGGCTTCTGCGAGCTGTGCGGCTACTCGCG
GGCCGAGGTGATGCAGCGACCCTGCACCTGCGACTTCCTGCACGGGCCGCGCACGCAGCGCCGAGCCGCAGCACAAATTGC
ACAAGCCCTACTAGGAGCAGAAGAACGGAAGGTGGAAATCGCCTTCTACCGGAAAGATGGGAGCTGCTTCCTATGTCTGGTGGA
TGTGGTGCCCGTGAAGAACGAGGATGGGGCTGTCATCATGTTCATCCTCAATTTTGAAGTAGTGATGGAGAAGGACATGGTGGG
GTCCCCGGCTCATGACACAAATCATCGAGGTCCTCCTACCAGCTGGCTGGCCCCAGGCCGCGCCAAGACCTTCCGCCTGAAGC
TGCCCGCGCTGCTGGCGCTGACGGCCCGGGAGTCGTCGGTGCGGTCGGGCGGCGCGGGCGGCGCGGGCGCCCCAGGAGC
AGTAGTAGTAGACGTAGATCTAACACCAGCGGCACCCAGCAGCGAGTCGCTGGCCCTGGACGAAGTGACAGCCATGGACAACC
ACGTGGCAGGACTAGGACCAGCAGAGGAACGACGCGCACTAGTAGGTCCCGGATCCCCACCACGAAGTGCACCAGGACAACT
ACCCTCACCACGAGCACATAGTCTTAATCCAGATGCATCAGGATCAAGTTGTAGTCTAGCACGAACACGATCACGCGAGAGTTGT
GCAAGTGTACGTCGTGCATCATCAGCAGATGATATAGAAGCTATGCGAGCTGGAGTGCTGCCCCCGCCACCGCGCCACGCCAG
CACCGGGGCCATGCACCCACTGCGCAGCGGCTTGCTCAACTCAACATCAGATTCCGACCTCGTGCGCTACCGCACCATTAGCA
AGATTCCCCAAATCACCCTCAACTTTGTGGACCTCAAGGGCGACCCCTTCTTGGCTTCGCCCACCAGTGACCGTGAGATCATAG
CACCTAAGATAAAGGAGCGAACCCACAATGTCACTGAGAAGGTCACCCAGGTCCTGTCCCTGGGCGCCGACGTGCTGCCTGAG
TACAAGCTGCAGGCACCGCGCATCCACCGCTGGACCATCCTGCATTACAGCCCCTTCAAGGCCGTGTGGGACTGGCTCATCCT
GCTGCTGGTCATCTACACGGCTGTCTTCACACCCTACTCGGCTGCCTTCCTGCTGAAGGAGACGGAAGAAGGCCCGCCTGCTA
CCGAGTGTGGCTACGCCTGCCAGCCGCTGGCTGTGGTGGACCTCATCGTGGACATCATGTTCATTGTGGACATCCTCATCAACT
TCCGCACCACCTACGTCAATGCCAACGAGGAGGTGGTCAGCCACCCCGGCCGCATCGCCGTCCACTACTTCAAGGGCTGGTTC
CTCATCGACATGGTGGCCGCCATCCCCTTCGACCTGCTCATCTTCGGCTCTGGCTCTGAGGAGCTGATCGGGCTGCTGAAGAC
TGCGCGGCTGCTGCGGCTGGTGCGCGTGGCGCGGAAGCTGGATCGCTACTCAGAGTACGGCGCGGCCGTGCTGTTCTTGCTC
ATGTGCACCTTTGCGCTCATCGCGCACTGGCTAGCCTGCATCTGGTACGCCATCGGCAACATGGAGCAGCCACACATGGACTCA
CGCATCGGCTGGCTGCACAACCTGGGCGACCAGATAGGCAAACCCTACAACAGCAGCGGCCTGGGCGGCCCCTCCATCAAGG
ACAAGTATGTGACGGCGCTCTACTTCACCTTCAGCAGCCTCACCAGTGTGGGCTTCGGCAACGTCTCTCCCAACACCAACTCAG
AGAAGATCTTCTCCATCTGCGTCATGCTCATTGGCTCCCTCATGTATGCTAGCATCTTCGGCAACGTGTCGGCCATCATCCAGCG
GCTGTACTCGGGCACAGCCCGCTACCACACACAGATGCTGCGGGTGCGGGAGTTCATCCGCTTCCACCAGATCCCCAATCCCC
TGCGCCAGCGCCTCGAGGAGTACTTCCAGCACGCCTGGTCCTACACCAACGGCATCGACATGAACGCGGTGCTGAAGGGCTTC
CCTGAGTGCCTGCAGGCTGACATCTGCCTGCACCTGAACCGCTCACTGCTGCAGCACTGCAAACCCTTCCGAGGGGCCACCAA
GGGCTGCCTTCGGGCCCTGGCCATGAAGTTCAAGACCACACATGCACCGCCAGGGGACACACTGGTGCATGCTGGGGACCTG
CTCACCGCCCTGTACTTCATCTCCCGGGGCTCCATCGAGATCCTGCGGGGCGACGTCGTCGTGGCCATCCTGGGGAAGAATGA
CATCTTTGGGGAGCCTCTGAACCTGTATGCAAGGCCTGGCAAGTCGAACGGGGATGTGCGGGCCCTCACCTACTGTGACCTAC
ACAAGATCCATCGGGACGACCTGCTGGAGGTGCTGGACATGTACCCTGAGTTCTCCGACCACTTCTGGTCCAGCCTGGAGATC
ACCTTCAACCTGCGAGATACCAACATGATCCCGGGCTCCCCCGGCAGTACGGAGTTAGAGGGTGGCTTCAGTCGGCAACGCAA
GCGCAAGTTGTCCTTCCGCAGGCGCACGGACAAGGACACGGAGCAGCCAGGGGAGGTGTCGGCCTTGGGGCCGGGCCGGG
CGGGGGCAGGGCCGAGTAGCCGGGGCCGGCCGGGGGGGCCGTGGGGGGAGAGCCCGTCCAGTGGCCCCTCCAGCCCTGA
GAGCAGTGAGGATGAGGGCCCAGGCCGCAGCTCCAGCCCCCTCCGCCTGGTGCCCTTCTCCAGCCCCAGGCCCCCCGGAGA
GCCGCCGGGTGGGGAGCCCCTGATGGAGGACTGCGAGAAGAGCAGCGACACTTGCAACCCCCTGTCAGGCGCCTTCTCAGG
AGTGTCCAACATTTTCAGCTTCTGGGGGGACAGTCGGGGCCGCCAGTACCAGGAGCTCCCTCGATGCCCCGCCCCCACCCCC
AGCCTCCTCAACATCCCCCTCTCCAGCCCGGGTCGGCGGCCCCGGGGCGACGTGGAGAGCAGGCTGGATGCCCTCCAGCGC
CAGCTCAACAGGCTGGAGACCCGGCTGAGTGCAGACATGGCCACTGTCCTGCAGCTGCTACAGAGGCAGATGACGCTGGTCC
CGCCCGCCTACAGTGCTGTGACCACCCCGGGGCCTGGCCCCACTTCCACATCCCCGCTGTTGCCCGTCAGCCCCCTCCCCAC
CCTCACCTTGGACTCGCTTTCTCAGGTTTCCCAGTTCATGGCGTGTGAGGAGCTGCCCCCGGGGGCCCCAGAGCTTCCCCAAG
AAGGCCCCACACGACGCCTCTCCCTACCGGGCCAGCTGGGGGCCCTCACCTCCCAGCCCCTGCACAGACACGGCTCGGACC
CGGGCAGTTAGTGGGGCTGCCCAGTGTGGACACGTGGCTCACCCAGGGATCAAGGCGCTGCTGGGCCGCTCCCCTTGGAGG
CCCTGCTCAGGAGGCCCTGACCGTGGAAGGGGAGAGGAACTCGAAAGCACAGCTCCTCCCCCAGCCCTTGGGACCATCTTCT
CCTGCAGTCCCCTGGGCCCCAGTGAGAGGGGCAGGGGCAGGGCCGGCAGTAGGTGGGGCCTGTGGTCCCCCCACTGCCCT
GAGGGCATTAGCTGGTCTAACTGCCCGGAGGCACCCGGCCCTGGGCCTTAGGCACCTCAAGGACTTTTCTGCTATTTACTGCTC
TTATTGTTAAGGATAATAATTAAGGATCATATGAATAATTAATGAAGATGCTGATGACTATGAATAATAAATAATTATCCTGAGGAGACT
CCAAAAAAAAAAAAAAAAAA

### hERG1aHN

ATGCCGGTGCGGAGGGGCCACGTCGCGCCACAAAACACATTCCTTGACACCATCATCCGCAAATTTGAGGGCCAGTCGCGCAA
GTTCATCATCGCCAACGCTCGGGTCGAGAACTGCGCTGTGATCTATTGCAATGACGGGTTCTGCGAGCTGTGCGGGTACTCGAG
AGCCGAGGTGATGCAGCGCCCATGCACCTGTGATTTCCTGCATGGCCCCCGGACACAGAGGCGCGCCGCCGCCCAGATCGCG
CAGGCCCTGCTGGGCGCCGAGGAGCGTAAGGTGGAGATTGCGTTTTACCGGAAGGATGGGTCCTGCTTCCTGTGCCTGGTCG
ACGTCGTGCCTGTGAAGAACGAGGACGGTGCCGTGATTATGTTCATCCTGAACTTCGAGGTGGTGATGGAGAAGGACATGGTG
GGCTCCCCCGCGCACGACACGAACCACCGAGGCCCTCCGACAAGCTGGCTGGCCCCTGGGCGCGCTAAGACCTTCCGCCTG
AAGCTCCCCGCCCTCCTCGCCCTGACGGCCCGGGAATCTAGTGTGCGCAGCGGCGGCGCGGGCGGCGCCGGTGCCCCGGG
TGCTGTGGTCGTCGACGTGGACCTGACCCCGGCTGCCCCCTCGAGCGAAAGTCTGGCCCTGGACGAGGTGACAGCGATGGAC
AACCACGTGGCCGGCCTAGGTCCTGCAGAGGAGAGGCGCGCGCTAGTGGGCCCTGGCTCCCCCCCCCGCTCCGCGCCGGGC
CAACTGCCGTCCCCCCGCGCCCACAGCCTGAACCCCGACGCCTCCGGCAGTAGCTGCAGCCTTGCCCGGACCCGGTCCCGC
GAATCTTGTGCGAGCGTCCGACGCGCCAGCAGCGCCGACGACATCGAAGCCATGCGCGCAGGCGTGCTGCCCCCCCCCCCC
CGACATGCGAGTACCGGCGCGATGCACCCCCTGCGGAGCGGCCTGCTCAACTCAACGTCTGATTCCGATTTGGTCAGGTACCG
GACAATTTCGAAGATCCCGCAGATCACCCTGAACTTCGTCGACCTCAAGGGCGACCCTTTCCTGGCCTCCCCCACCAGCGACC
GCGAGATCATAGCACCGAAGATCAAGGAGCGGACCCACAACGTGACAGAGAAGGTGACCCAGGTCCTGAGTCTCGGCGCCGA
TGTGTTGCCTGAATATAAGCTGCAAGCTCCCCGGATCCACAGGTGGACGATCCTGCACTACAGCCCGTTCAAGGCCGTCTGGGA
CTGGCTCATCCTGCTCCTGGTGATCTACACAGCCGTGTTTACCCCCTACTCCGCCGCCTTTCTGCTGAAGGAGACCGAAGAGGG
CCCCCCTGCGACCGAGTGCGGCTACGCCTGCCAACCCCTGGCCGTGGTGGACCTGATCGTGGACATCATGTTCATCGTGGACA
TCCTAATCAACTTCCGTACGACGTACGTCAACGCCAACGAGGAGGTCGTCAGCCATCCAGGCCGGATCGCCGTGCACTACTTCA
AAGGCTGGTTCCTGATCGACATGGTGGCCGCAATCCCATTCGACCTCCTGATATTCGGCTCCGGGTCCGAGGAGTTGATCGGCC
TGTTGAAAACCGCCCGCCTCCTCCGACTGGTCCGTGTCGCCCGCAAGCTGGACCGCTACTCCGAGTACGGCGCCGCAGTGTT
GTTCCTGCTGATGTGCACCTTCGCCCTCATCGCTCACTGGCTCGCCTGCATCTGGTACGCGATTGGGAACATGGAGCAGCCCCA
CATGGATAGTAGGATCGGGTGGCTCCACAACCTGGGGGACCAGATTGGGAAACCGTACAACAGCAGCGGCCTGGGCGGCCCC
TCCATCAAGGACAAGTACGTGACCGCACTATACTTCACTTTCAGTAGTCTGACTAGCGTGGGCTTCGGCAATGTGAGCCCCAATA
CCAATAGCGAAAAGATCTTCAGCATCTGCGTGATGCTCATCGGTTCCCTGATGTACGCGTCGATCTTCGGGAACGTTAGCGCCAT
CATTCAGCGCCTGTACAGCGGCACAGCACGGTACCACACCCAGATGCTCCGGGTCCGAGAGTTCATACGGTTCCACCAGATCC
CCAACCCCCTACGACAGCGCCTCGAGGAGTACTTCCAGCACGCCTGGTCTTACACGAATGGCATCGACATGAACGCTGTCTTGA
AGGGCTTCCCCGAGTGCCTCCAAGCCGATATCTGCCTACATCTGAACCGCTCCCTCCTGCAGCACTGCAAGCCATTCCGAGGG
GCGACCAAGGGGTGCCTGCGGGCTCTGGCAATGAAGTTCAAGACGACGCACGCCCCCCCAGGCGACACACTGGTGCATGCC
GGCGACCTCCTGACTGCTCTGTACTTCATCTCACGGGGCTCAATCGAGATCCTTCGTGGCGATGTGGTGGTGGCGATCCTGGG
CAAGAATGACATTTTCGGCGAGCCCCTGAACCTGTACGCCCGACCCGGCAAGTCGAACGGCGACGTGAGGGCCCTGACGTACT
GCGACCTGCACAAGATCCACCGGGACGACCTGCTCGAGGTCCTGGATATGTACCCCGAGTTTAGCGACCACTTCTGGAGTAGC
TTAGAGATAACGTTCAACCTGCGAGACACCAACATGATCCCCGGTTCGCCCGGCTCGACCGAGTTAGAAGGGGGCTTTTCCCGA
CAGAGAAAGCGCAAGCTGAGCTTCCGCCGCCGCACCGACAAGGACACAGAGCAGCCAGGGGAGGTGAGCGCCCTGGGCCCC
GGCCGAGCAGGGGCAGGCCCGTCCAGTCGGGGGCGCCCTGGGGGCCCGTGGGGTGAGTCACCGTCAAGCGGCCCGTCCAG
TCCCGAGAGTAGTGAAGACGAGGGCCCCGGTCGAAGTAGTTCCCCCCTGCGCCTGGTGCCGTTCTCGAGTCCGCGCCCCCCT
GGCGAGCCCCCCGGCGGGGAGCCCCTGATGGAGGATTGTGAGAAGAGTAGCGACACGTGTAACCCCCTGAGCGGCGCCTTCA
GTGGAGTGAGCAATATCTTCTCGTTCTGGGGGGACTCCCGCGGACGGCAGTACCAGGAGCTTCCCCGATGCCCCGCACCCACT
CCCAGCCTCCTGAATATCCCCCTGTCCTCACCCGGTCGGCGACCACGCGGAGACGTGGAGAGTCGCCTGGACGCCCTGCAGA
GGCAGCTGAACCGCCTGGAGACCCGCCTGAGTGCCGACATGGCCACCGTCCTGCAGCTCCTCCAACGGCAGATGACCCTGGT
CCCCCCCGCCTACTCCGCCGTGACGACCCCTGGCCCCGGCCCGACCTCGACCAGCCCTCTATTGCCTGTTAGCCCCCTGCCTA
CCCTCACCCTCGATAGCCTGTCCCAGGTCTCCCAGTTCATGGCGTGTGAAGAGTTGCCCCCCGGCGCTCCAGAGCTGCCCCAG
GAAGGCCCGACCCGACGTCTCTCTCTACCCGGCCAGCTCGGCGCCCTGACCTCCCAGCCCCTGCACAGACACGGAAGCGACC
CTGGCTCGTAGTGGGGCTGCCCAGTGTGGACACGTGGCTCACCCAGGGATCAAGGCGCTGCTGGGCCGCTCCCCTTGGAGG
CCCTGCTCAGGAGGCCCTGACCGTGGAAGGGGAGAGGAACTCGAAAGCACAGCTCCTCCCCCAGCCCTTGGGACCATCTTCT
CCTGCAGTCCCCTGGGCCCCAGTGAGAGGGGCAGGGGCAGGGCCGGCAGTAGGTGGGGCCTGTGGTCCCCCCACTGCCCT
GAGGGCATTAGCTGGTCTAACTGCCCGGAGGCACCCGGCCCTGGGCCTTAGGCACCTCAAGGACTTTTCTGCTATTTACTGCTC
TTATTGTTAAGGATAATAATTAAGGATCATATGAATAATTAATGAAGATGCTGATGACTATGAATAATAAATAATTATCCTGAGGAGACT
CCAAAAAAAAAAAAAAAAAA

**Supplementary Video 1. *hERG1a* and *1b* mRNAs form dynamic heterotypic condensates in HeLa cells.** Fluorescently labeled *hERG1a* (red) and *1b* (cyan) mRNAs were co-transfected into HeLa cells and imaged 24 hours later. Co-transfected mRNAs formed heterotypic condensates, appearing as white, that move inside the cell and undergo multiple fission and fusion events.
